# *alpseq*: an open-source workflow to turbocharge nanobody discovery with high-throughput sequencing

**DOI:** 10.1101/2025.10.22.661623

**Authors:** Kathleen Zeglinski, Jakob Schuster, Jaison D Sa, Amy Adair, Jing Deng, Phillip Pymm, Matthew E. Ritchie, Rory Bowden, Wai-Hong Tham, Quentin Gouil

## Abstract

Nanobodies have emerged as promising tools for many biotechnological applications due to their small size, high stability, and remarkable binding specificity. Next-Generation Sequencing (NGS) enables deep profiling of large nanobody libraries and panning campaigns, however the scale and diversity of nanobody NGS datasets presents a significant bioinformatic challenge. To this end, we have developed *alpseq*, an optimised, open-source software pipeline designed specifically for the efficient and accurate processing of NGS data from nanobody libraries and panning campaigns. *alpseq* is also paired with a PCR-free sequencing library preparation protocol to allow researchers to easily generate their own data while avoiding biases. The *alpseq* software pipeline is composed of two parts: a pre-processing module written in Nextflow efficiently handles raw nanobody reads in a single line of code. These results are then fed into the analysis module, which contains a comprehensive suite of functions for quality control, diversity analysis, identification of enriched sequences and clustering. *alpseq* also creates a user-friendly interactive report which empowers scientists to explore their data without the need for extensive bioinformatic experience. Sophisticated panning campaign designs are supported, such as replicates and comparisons between different pans to find cross-binding leads. *alpseq* thus generates insights into the nanobody selection process and delivers a list of lead candidates for further experimental validation and downstream applications. *alspeq* is available at https://github.com/kzeglinski/alpseq.

## Introduction

Nanobodies, also known as single-domain antibodies, are the variable antigen-binding (VHH) regions from camelid heavy-chain-only antibodies (1). First discovered in 1989 in dromedary camels, heavy-chain-only antibodies are able to recognise a wide variety of antigens despite their smaller size (2). They have since been discovered in other camelids, including llamas and alpacas (3) as well as some species of cartilaginous fish, notably sharks (4). Irrespective of the species from which they are derived, the nanobodies’ smaller size allows them to target obstructed epitopes, better penetrate tissues and even cross the blood-brain barrier (1). Nanobodies also have higher thermal stability and solubility than conventional monoclonal antibodies, as well as lower immunogenicity, making them attractive for clinical development (1).

Target-specific nanobodies are typically selected from libraries using display technologies such as phage display (5), yeast or ribosome display (6). These libraries are often derived from immunised animals, but may be also be naïve (from non-immunised animals) or synthetic (6). The selection process, known as ‘biopanning’ involves multiple rounds of selection against a target antigen, with non-binders being removed via washing to enrich for target-specific nanobodies. Following biopanning, the typical next step is to perform low-throughput Sanger sequencing of the final round output (6). This does not scale well and only captures a few hundred of the potentially tens of thousands of unique clones that remain after the panning process (7). Thus, in the field of antibody discovery, there has long been interest in coupling highthroughput, next-generation sequencing (NGS) with display technologies in order to capture a more diverse set of potential binders (7–9). Nanobodies, given their shorter length (∼400 bp) and single chain, are even more suitable for NGS than antibodies (>800 bp) as they can be sequenced in their entirety with paired 2 *×* 300 bp reads. However until recently, NGS was used only ‘sporadically’ (6) in nanobody discovery (10–12), with a recent influx of publications suggesting that it is beginning to be more widely adopted by the field (13–16). Nevertheless, further optimisations of the sequencing method would likely increase the adoption of NGS in nanobody discovery. Currently, most studies rely on PCR-based approaches for sequencing library preparation (10, 16) which is known to introduce bias. Although the bias can be corrected bioinformatically through the use of unique molecular identifiers (UMIs) (17), a PCR-free approach would be more straightforward, eliminating the bias entirely.

Additionally, we believe a significant factor in the slow uptake of NGS by the nanobody field is the limited availability of open-source and user-friendly software with which to analyse the resulting data. Although several published studies share their analysis code, it is specific to a particular dataset or selection approach (10, 15) and bioinformatic expertise would be required in order to replicate the results. Several commercial solutions exist (18–20) but the costs associated with these can be prohibitive and their closed-source nature can make it difficult to tailor analysis to specific projects. As nanobodies in their basic structure are similar to antibody heavy chain domains (1), it is often possible to use antibody analysis pipelines to process nanobody data, although there are some tools that require both a heavy and light chain (21, 22). Yet even in the larger field of antibody discovery, there are still few open source pipelines available, many of which are free for academic use only (23), specialise in dealing with repertoire-derived data rather than *in vitro* selection data (24, 25) or do not handle the full end-to-end analysis (from raw sequencing reads to top candidates)(26).

Open-source software has an important role in all forms of scientific data analysis, allowing researchers to freely explore their data and bioinformaticians to save time by tailoring existing code to suit their needs (27). When coupled with clearly documented laboratory protocols, open-source software has the power to democratise entire fields of research and drive innovation. To this end, we propose *alpseq*: a freely available end-to-end workflow that empowers researchers to routinely generate and analyse nanobody NGS data in-house. The insights gained through this analysis have the potential to accelerate nanobody discovery by identifying a more diverse range of sequences, and by generating large, pre-processed datasets that can be fed into various artificial intelligence algorithms.

## Results

### The *alpseq* sequencing workflow reliably generates large volumes of nanobody data

The PCR-free *alpseq* library preparation protocol is outlined in Figure 1A, with further details in the Materials and Methods section (see below). Across >10 sequencing runs it has generated large volumes of data, with an average of 10 million full-length nanobody sequences per MiSeq run and 75 million per NextSeq 2000 P1 run (Figure 1B), after accounting for pairs of reads that do not merge and the phiX spike-in. Read quality in the NextSeq 2000 runs represented a significant improvement over MiSeq, particularly towards the end of read 2 (Figure 1C).

**Fig. 1.**
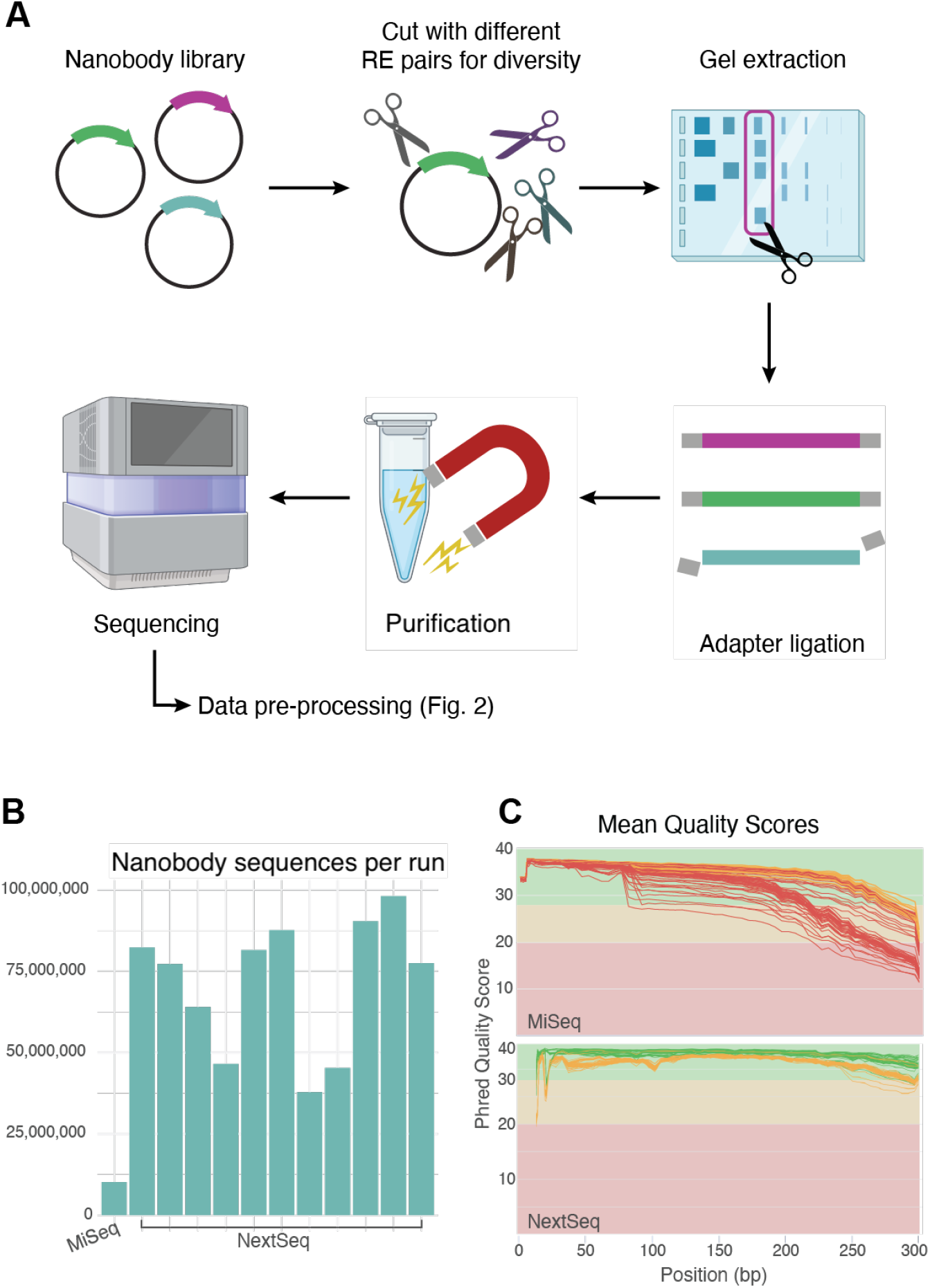
Overview of the *alpseq* laboratory workflow. **(A)** Steps involved in the *alpseq* laboratory workflow. Nanobody domain sequences are cut from nanobody library plasmids using different pairs of restriction enzymes (RE) to maintain sequence diversity near the sequencing adapters. Gel extraction is used to isolate the nanobody sequences, which are then purified with magnetic beads before having adapters ligated. Sequencing is finally performed using the NextSeq 2000 instrument with the P1 2*×*300 cycles kit. **(B)** Bar chart showing the number of reads generated across 12 *alpseq* runs, 1 on MiSeq and the rest on the NextSeq 2000. **(C)** FastQC mean quality score plots from representative *alpseq* runs using the Miseq (top) and NextSeq (bottom).

### Fast and efficient nanobody data pre-processing with *alpseq*’s Nextflow pipeline

Given the large volumes of data generated by *alpseq* (in contrast to Sanger sequencing), it was necessary to develop an efficient bioinformatic pipeline to pre-process the data prior to analysis of clone enrichment. The steps of this pipeline are outlined in Figure 2, and involve trimming and merging of reads to generate a contiguous nanobody sequence, before annotating them to identify their germline genes and CDRs. Quality control is performed throughout the pipeline, and collated in a comprehensive QC report (example report available on github). *alpseq* is written in the Nextflow workflow language (28), which efficiently parallelises the analysis of many samples (Figure 2B) at once and allows the entire pipeline to be executed with a single line of code. Nextflow also offers a graphical user interface through the Seqera Platform (Supplementary Figure 1), facilitating the uptake of *alpseq* by users without extensive bioin-formatics experience.

**Fig. 2.**
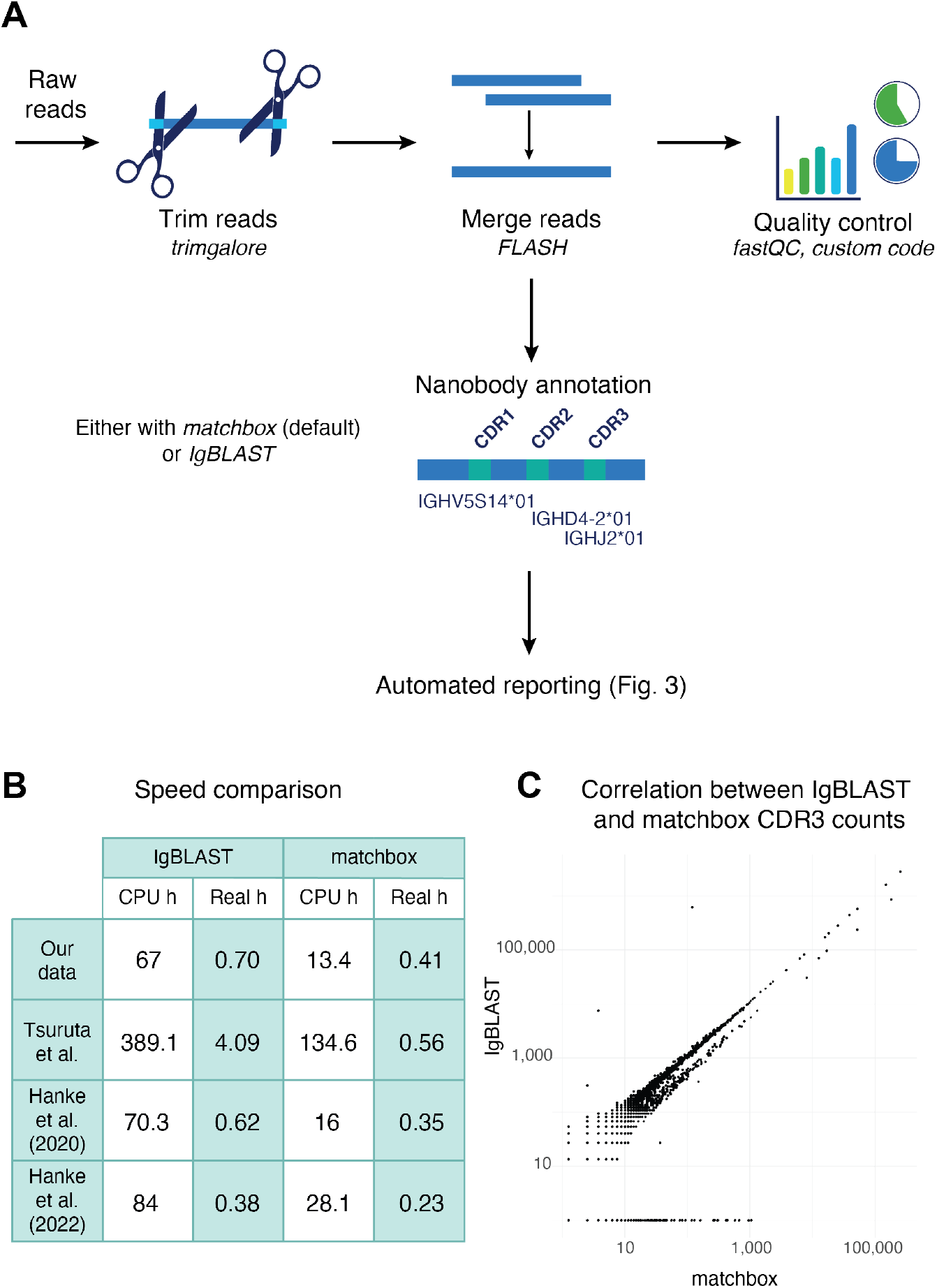
Overview of the *alpseq* bioinformatics pipeline. **(A)** In the pre-processing step, paired sequencing reads are trimmed, merged and annotated to identify the nanobody sequences and their key features. Quality control of the sequencing is also performed. Finally, in the report generation step, users with bioinformatic experience can perform a manual analysis using the *alpseq* R package, or an interactive HTML report can be automatically generated using the provided sample sheet. **(B)** Performance of *alpseq* on the data provided with this study (ENA PRJEB90877), and three public datasets (10, 31, 32) in both CPU hours (no parallelisation) and real hours (with parallelisation) when using either IgBLAST or matchbox for nanobody annotation. **(C)** Comparison of clone counts generated from the Hanke et al. (2020) (10) dataset using IgBLAST vs *matchbox*.

Despite parallelisation with Nextflow, the nanobody annotation step remains computationally intensive when performed with the gold-standard IgBLAST software (29), requiring hundreds of CPU hours for a single NextSeq run (Figure 2B). This limits the deployment of *alpseq* with IgBLAST annotation to high-performance or cloud computing systems, which can be expensive and difficult to access. To solve this issue, *alpseq* also implements a new annotation strategy using the recently released *matchbox* software (30) (https://github.com/jakob-schuster/matchbox). *matchbox* annotates the V gene and CDR3 of nanobodies significantly faster than IgBLAST (exact speedup depends on the size of the dataset, typically at least 2–5x; Figure 2B), while also allowing mismatches from mutations and indels, with the results being highly concordant (97.85% agreement, *r* = 0.9425; see Figure 2C). Given the speed of *matchbox*, and that the choice of annotation software has no effect on the selection of a top 100 set of sequences, *matchbox* is implemented as the default annotation method in *alpseq*. This makes analysis on personal computers possible.

### Automated interactive reporting with *alpseq* facilitates discovery of enriched nanobody sequences

Following pre-processing, *alpseq* can be used to perform enrichment analyses and reporting in two different ways: automated or manual. In the automated mode, interactive HTML reports that can be opened in any web browser are automatically generated by the *alpseq* Nextflow pipeline following template analyses specified from the samplesheet. The manual mode allows for more custom analysis via an accompanying R package (*alpseqR*) containing the comprehensive set of functions that underlie *alpseq*’s reports.

*alpseq* analyses nanobody sequences at the CDR3 level, and determines their enrichment by calculating their log_2_ fold change (logFC) across the panning process (see Materials and Methods). CDR3s are clustered using CD-HIT (33) to minimise redundancy when selecting top candidates to take forward for functional validation. Clusters with less than 100 counts per million (CPM) at the end of the panning process are discarded, as empirical validation suggests that these are likely false positive hits (Supplementary Figure 2A). Otherwise, all information about clusters is retained, as once binding data is available it allows the NGS data to be mined for related clones that may have better binding affinity or developability. These clusters and their associated enrichment or abundance values are presented in interactive MDS plots and cladograms (Figure 3) to visualise the overall sequence diversity landscape (which can reveal ‘super-clusters’ of nanobodies that may bind different epitopes of the target). The size of the dots in the MDS plot and cladogram further indicate the abundance of the enriched clones composing each cluster in the final round. The visualisations are linked to a table that can be filtered/sorted within the report, or downloaded for further external analysis (Figure 3B). With these interactive features, users are empowered to select their own favourite candidates from the NGS results, without requiring bioinformatic knowledge.

**Fig. 3.**
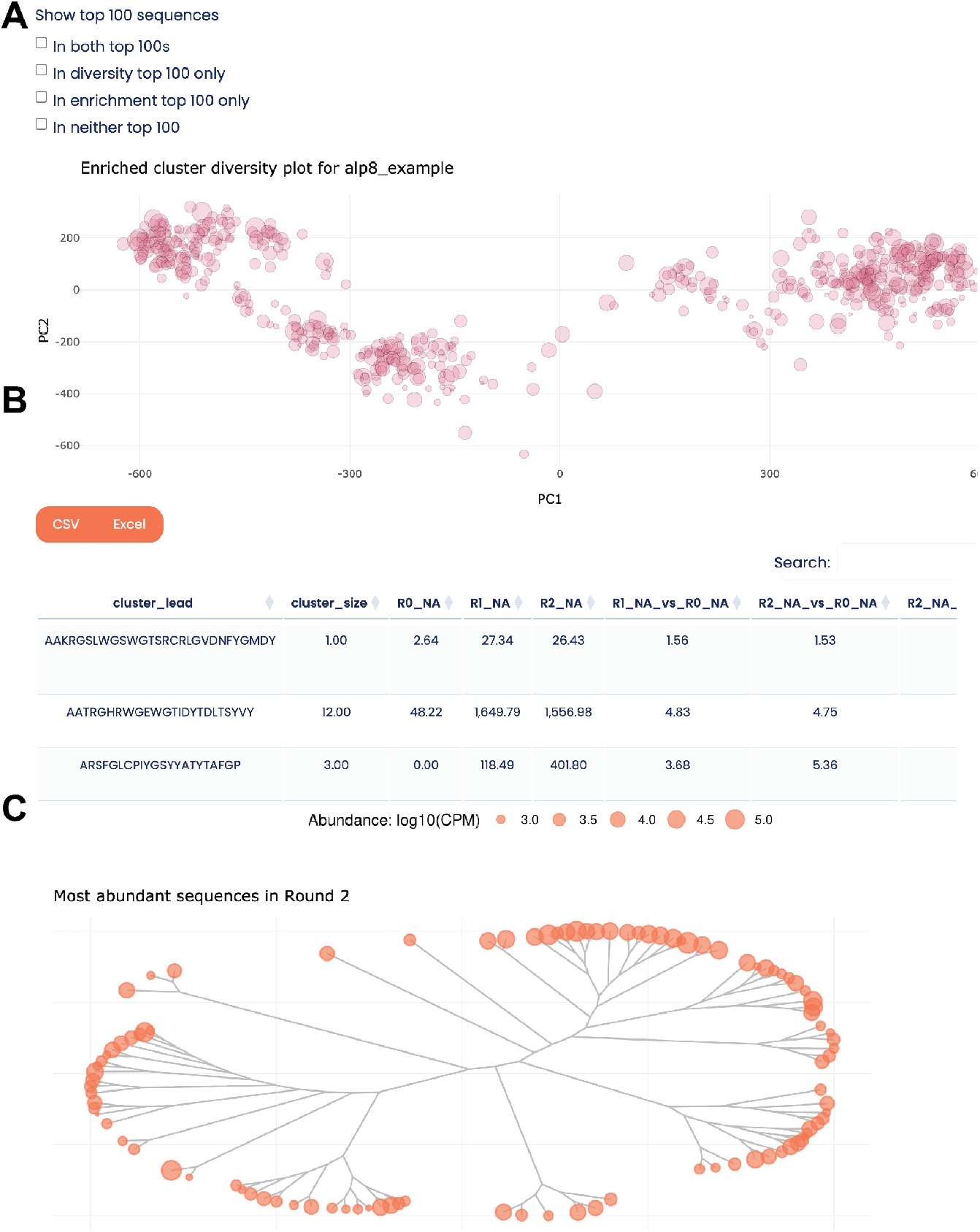
Functionality for selecting candidate binders. **(A)** An interactive MDS plot is used to visualise the sequence diversity landscape, revealing ‘superclusters’ of nanobodies that likely target the same/similar epitopes. **(B)** The plot is dymamically linked to a table that provides detailed information about the enrichment and abundance of each nanobody cluster throughout the biopanning process. **(C)** Cladogram visualisation of the diversity of the most abundant nanobody sequences in a given round.

As there can be many hundreds of enriched nanobody clusters, it is often necessary to narrow these down further to a top 100 that can be validated in the lab. Although these can be selected manually by researchers as described above, *alpseq* also provides a list of 100 nanobodies that are likely to be good candidates for target-binding. Highly abundant (CPM > 1000) and highly enriched (logFC > 2.5) clusters are prioritised to fill this list, as ELISA and BLI data suggests that there are very few non-binders above these cutoffs (Supplementary Figure 2). The most abundant clones are chosen from each cluster. Typically, there are not 100 nanobody clusters in a dataset that meet these thresholds, so the remaining picks are taken from any enriched cluster and failing that, additional picks are made from within already-selected clusters.

### *alpseq* effectively handles complex study designs

To highlight *alpseq*’s analysis capabilities, we describe two more advanced use cases: the handling of replicates and the comparisons of different pans.

### Replicates

The enrichment of a particular sequence across multiple replicate pans has been shown to increase the robustness of candidate selection in peptide screens, given the large variability in phage display (34). Thus, *alpseq* supports the selection of top candidates from a set of replicate pans, by prioritising nanobodies that are enriched in multiple replicates when choosing the set of top 100 sequences. Replicate information is also present in all output, allowing users to see which replicate(s) a particular cluster appears in. For all pans containing replicates, a series of all-vs-all scatterplots comparing the normalised abundance of nanobodies is generated (Figure 4A). This can help to identify abnormal replicates that may be the result of an issue in the biopanning, mislabelled samples or other problems in the wet lab, which informs the selection of candidate nanobodies.

**Fig. 4.**
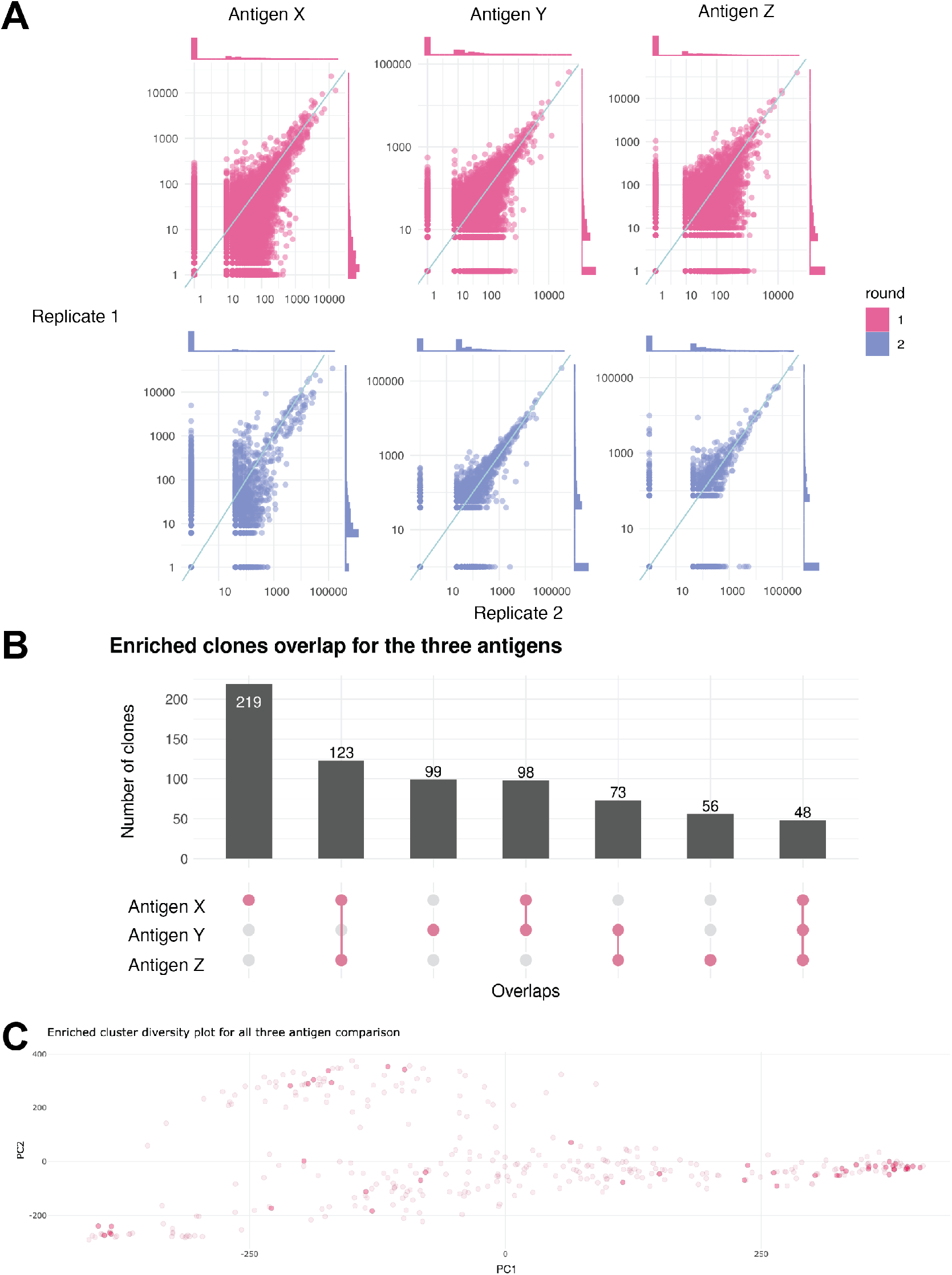
*alpseq* supports complex study designs. **(A)** Scatterplots comparing the normalised abundance (CPM) of nanobodies across replicates. Density plots along the top and right sides show the distributions of nanobody abundance in each replicate. The *y* = *x* line is annotated on all plots. **(B)** An UpSet plot showing the overlap of enriched nanobody sequence clusters in three pans against three different antigens. The pink dots along the x axis represent the overlaps, and the bar chart shows how many nanobody clusters belong to each overlapping category. **(C)** A variant on the interactive MDS plot and table that indicates which pan/antigen each cluster is enriched for.

### Comparing pans

*alpseq* can also be used to analyse nanobodies across multiple pans. For a single antigen, this could be to compare different panning conditions (different ways of preparing the antigen, or different display techniques). Comparing pans across multiple antigens might be used to find cross-reactive binders (e.g. nanobodies recognising multiple viral strains) or to subtract a set of enriched sequences (i.e. to find nanobodies that bind antigen x but not y). This is done by first clustering all nanobodies enriched in all pans to be compared, to account for the fact that highly similar (but not identical) clones may become enriched in different pans even if they bind in a similar way. Then, the overlap of clusters between pans is examined, and visualised in an UpSet plot (35), as well as a linked MDS plot and table, colour-coded and labelled to show which pan(s) each enriched nanobody cluster belongs to (Figure 4B).

### Demonstration of *alpseq*’s capability on public data

To demonstrate the robustness of the *alpseq* software pipeline to different sample preparation strategies, we ran *alpseq* on the publicly available data from Hanke *et al*. 2020 (10). *alpseq* was able to perform a complete analysis, from raw reads to interactive report in 30 minutes (using IgBLAST) or 12 minutes (using matchbox), with a single line of code. The Ty1 clone identified by Hanke *et al*. was also identified by *alpseq*, and featured in the list of top 100 sequences (Supplementary Figure 3). The interactive report generated from this data is available on github. 8 other highly similar nanobody sequences were clustered together with Ty1 by *alpseq*, and approximately 300 similar nanobody clusters were grouped together into a ‘super cluster’ on the MDS plot, suggesting that they may bind similar epitopes (Supplementary Figure 3). To further demonstrate the generalisability of *alpseq*, we also ran the pipeline on two additional public datasets from Tsuruta *et al*. (31) and Hanke *et al*. 2022 (32). *alpseq* was able to efficiently process these datasets (Figure 2B), visualise the sequence diversity landscape of the same library panned with different antigens in the Tsuruta *et al*. dataset (Supplementary Figure 4A) as well as compare the different pans in the Hanke *et al*. 2022 dataset (Supplementary Figure 4B).

## Discussion

Given the increasingly important roles of nanobodies in research and medicine (1), streamlining their discovery process has been a long-standing goal for the field (12–14). Highthroughput sequencing can achieve this by rapidly and deeply profiling phage, yeast or ribosome display outputs, thereby generating large datasets that are able to be mined for rare candidates or fed into machine learning algorithms for further insights. However, a lack of freely-available and easy to use tools with which to process this data has limited the uptake of NGS in the nanobody community (24–26). To address this gap, we have developed *alpseq*, an end-to-end wet- and dry-lab workflow that reliably generates large volumes of nanobody sequencing data, and processes it in a computationally efficient way. This is achieved through the use of the nextflow programming language and the *matchbox* software for nanobody annotation, which both deliver significant speedups (Figure 2B). However, it is important to note that the *matchbox* annotation is specifically designed for use with enriched nanobody panning data (see methods for the specific assumptions it makes) and therefore would not be suitable as a general-purpose nanobody annotation tool. *alpseq* is designed to be straightforward to use, running in a single line of code, and generates interactive reports that can be used by researchers without computational skills to explore their data and choose top candidates for further experimental validation. To date, *alpseq* has been used to process hundreds of internal datasets (36, 37).

The support for analysis of complex experimental designs in *alpseq* lowers the barrier to using sophisticated and replicated selection strategies. By streamlining data generation, processing and interpretation, *alpseq* allows researchers to focus on higher-level experimental insights rather than having to spend time writing low-level data processing code. This not only enhances reproducibility but also accelerates nanobody discovery.

Beyond its automated capabilities, the *alpseq* R package provides a flexible interface that bioinformaticians can use to perform their own custom analyses. For example, all of the thresholds for abundance, enrichment and sequencing identity are tunable by the user, allowing them to chose values that are suitable for their dataset (e.g. if many unique clones remain at the end of panning, stricter thresholds can be set and vice versa if the sample is highly clonally restricted). Thus, the *alpseq* workflow can easily be applied to sequence and analyse the outputs of other display technologies, and the computationally efficient pre-processing module can be applied to any nanobody sequencing data, including repertoire sequencing. Pre-processed data from *alpseq* can also be used to train or run machine learning models for predicting developability (ideal drug properties), structure and even the residues involved in binding (38–41). Because *alpseq* is open-source, there is the flexibility to add these tools as additional steps in the pipeline, allowing *alpseq* to continue to evolve and leverage the latest improvements in the bioinformatics and nanobody fields.

Although designed primarily for high-throughput sequencing data, the *alpseq* bioinformatics pipeline can also be used to process Sanger-derived nanobody sequences through its ‘single-read’ mode, allowing it to integrate seamlessly into existing workflows. This mode also allows *alpseq* to process data from long-read sequencing platforms.

In summary, *alpseq* is a comprehensive sample preparation and bioinformatic workflow that enables high-throughput sequencing of nanobody biopanning samples. *alpseq* is designed to be easy-to-use, and is freely available so as to facilitate the wider uptake of NGS in the nanobody community. We hope that by making the generation and processing of large nanobody sequencing datasets accessible, that not only will it be easier for researchers to discover a wider range of nanobodies from their panning experiments, but also that a more open culture of data-sharing might develop which would benefit the entire research community.

## Materials and Methods

### NGS library preparation and sequencing

Plasmids from phage libraries were digested and purified by gel extraction to isolate the nanobody domain sequences. Three different pairs of restriction enzymes were used to maintain sequence diversity at both ends, which is critical for Illumina sequencing with the 2-colour chemistry (42). Paired-end sequencing libraries were prepared using the NEBNext Multiplex Oligos for Illumina (Cat # E7395) in a PCR-free manner according to the manufacturer’s instructions. The samples were size-selected using AMPure XP beads with a 0.7*×* ratio (Cat # A63881) and quantified by qPCR using the KAPA Library Quantification Kit (Cat # KK4873) and sequenced on an Illumina NextSeq 2000 instrument with the P1 2 *×* 300 cycle kit (with latest XLEAP-SBS chemistry) and a 20% PhiX spikein.

### Pre-processing of nanobody NGS reads

Raw sequencing reads were trimmed to remove sequencing adapters with TrimGalore (v0.6.7) (43) and then merged using FLASH (v1.2.11) (44) with a maximum overlap of 250 bp to avoid carrying adapter dimers through to the next step. Annotation of nanobody sequences was performed either using Ig-BLAST (v1.19.0) (29) or using *matchbox* (v0.1) (30). The default tool for nanobody annotation in *alpseq* is matchbox, although this can be changed using the ‘–use_igblast’ parameter. A detailed description of how *matchbox* was used for nanobody annotation can be found below. The default reference database included in alpseq is for alpaca (*Vicugna pacos*) built from IMGT (45), although users can supply their own if desired. In R (v4.4.1), nanobodies were collapsed into clones at the CDR3 level, and CDR3s with a count of 1 were removed as these likely result from sequencing errors. To allow for comparison between pans, the counts of clones were then normalised through conversion to counts per million (CPM). The entire pre-processing pipeline is automated using Nextflow (28), and can be run with a single line of code.

### Annotation with *matchbox*

First, a sample of 1000 random reads is taken, and processed to determine the 15 most common V genes used in these nanobodies. This is an optimisation made with the assumption that enriched nanobody libraries will contain relatively few V genes. The number of V genes taken forward can be tuned with the ‘– num_v_genes’ parameter (if users expect that their library will be more diverse), and the matching is done to V gene reference sequences, with a 20% error rate to allow for sequencing errors and any mutations/deviation from the reference sequence. These 15 most common V genes are then trimmed to just before the last conserved cysteine. Then, the sequence between the FWR4 and the codon after the V gene (i.e. the conserved cysteine) is labelled as the CDR3 (FWR4 and V gene are matched with an error rate of 30% by default, but this can be tuned with the ‘–mb_error_rate’ parameter). The whole nanobody sequence and CDR3 are then translated, and the output written to a CSV. Checks for stop codons and frameshifts are done upon reading this CSV into R, specifically we check whether stop codons exist in the amino acid sequence and whether the CDR3 is a substring of the whole amino acid sequence (to ensure they are in the same reading frame). If either of these conditions are not met, the sequence is termed ‘unproductive’.

### Determination of enriched and top 100 clones

Enrichment of clones with a count > 1 at the end of panning was measured using the log_2_ fold change (logFC), calculated as

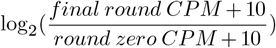

10 was added to the numerator and denominator to minimise the impact of lowly abundant clones on the analysis as their counts may not be reliable. In this particular analysis, clones with a logFC of at least 1.5 were considered enriched (although this is tunable). In order to minimise redundancy, enriched clones were then clustered by CDR3 amino acid sequences using CD-HIT (default length and identity threshold is 80%, but this can be tuned by the user) (33) and those clusters with less than 100 CPM at the end of panning were removed (as in our experience despite being enriched according to the logFC threshold, these nanobodies rarely bind the target by either ELISA or BLI, see Supplementary Figure 2). To visualise the sequence diversity landscape of enriched clones, a representative CDR3 from each cluster was chosen based on abundance. These CDR3s were then subjected to an all-vs-all pairwise alignment using the pwalign (46) (v1.4.0) R package with a gap opening penalty of -10 and a gap extension penalty of -1. The BLOSUM62 substitution matrix was used to try and account for similar chemical properties of certain amino acids. The result of this all-vs-all alignment (a distance matrix, where each element is a score measuring the similarity of two CDR3 sequences) was then used to perform MDS plot using the prcomp_irlba function from the irlba R package (47) (v2.3.5.1). The result of this was used for the interactive MDS plots and cladograms.

To find a set of top 100 sequences for further testing, the most abundant representative from each cluster was selected, prioritising clusters with logFC > 2.5 or CPM > 1000 or that consistently increase in abundance across the rounds of panning. If this does not produce 100 unique sequences, additional picks are made from selected clusters until a top 100 can be filled. Custom selections of clones of interest are also possible, using either enrichment or abundance or both to filter/rank selections.

### Overlap and subtraction

When comparing different pans, all enriched sequences from all pans are pooled and reclustered to account for similar but not identical clones being shared across pans. These new clusters are then reported as overlapping (i.e. cross-reactive) or removed (in the case of subtraction).

## ACKNOWLEDGEMENTS

We thank Daniel Brown and Jafar Jabbari for suggesting the PCR-free library preparation approach. We thank Stephen Wilcox and Sarah MacRaild from WEHI Genomics platform for their assistance in sequencing the nanobody libraries. We thank the WEHI Research Computing Platform for providing access to Seqera Cloud.

## Data availability statement

The example sequencing data highlighted in this study is available at the ENA under accession PRJEB90877. The publicly available data from Hanke *et al*. (10) is available from the SRA, under BioProject ID PRJNA638614.

## Author contributions

Conceptualisation: KZ, QG, MER, WHT; Data acquisition: JDS, AA, JD, PP; Data analysis: KZ; Software: KZ, JS; Interpretation: KZ, JDS, WHT, PP, QG; Resources: MER, WHT, QG, RB Drafting of manuscript and figures: KZ; Review and editing: KZ, MER, WHT, QG. All authors read and approved the final manuscript.

