## Supplementary for "*alpseq*: an open-source workflow to turbocharge nanobody discovery with high-throughput sequencing"

### Supplementary Note 1: Running *alpseq* through the Seqera Platform GUI

Seqera offers a graphical interface for running nextflow pipelines using the Seqera Platform (https://seqera.io/platform/). *alpseq* is designed to be compatible with this, allowing those with limited bioinformatic experience to run the program from their web browser, without the need to use the command-line.

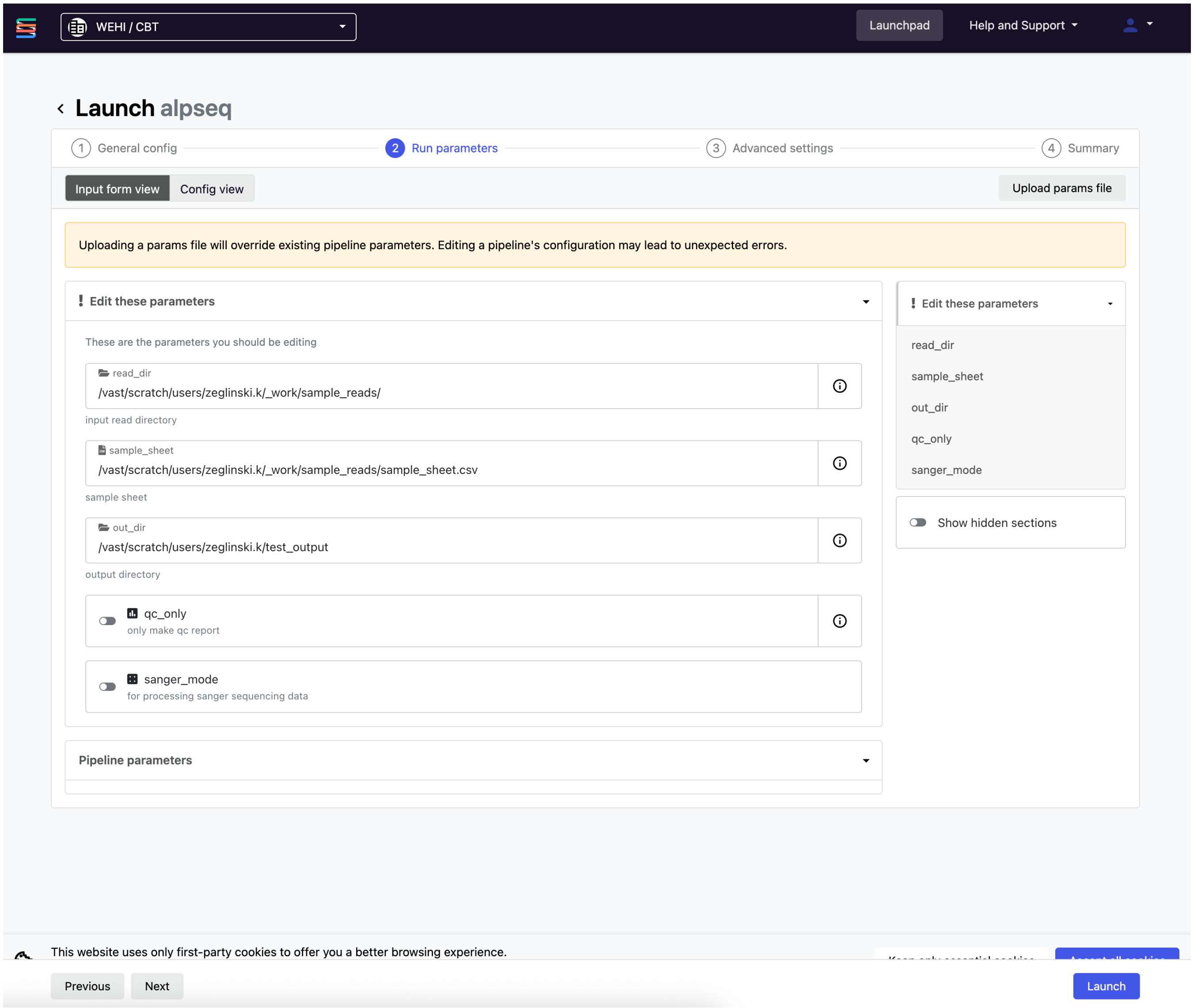


**Fig. 1.** Screenshot showing the graphical interface for *alpseq*, provided through the Seqera Platform.

### Supplementary Note 2: Empirical selection of abundance and enrichment cutoffs

Given that phage display is known to generate false positive hits (34), we used antigen binding status (as determined by ELISA and BLI) to try to empirically determine appropriate cutoffs for abundance and enrichment.

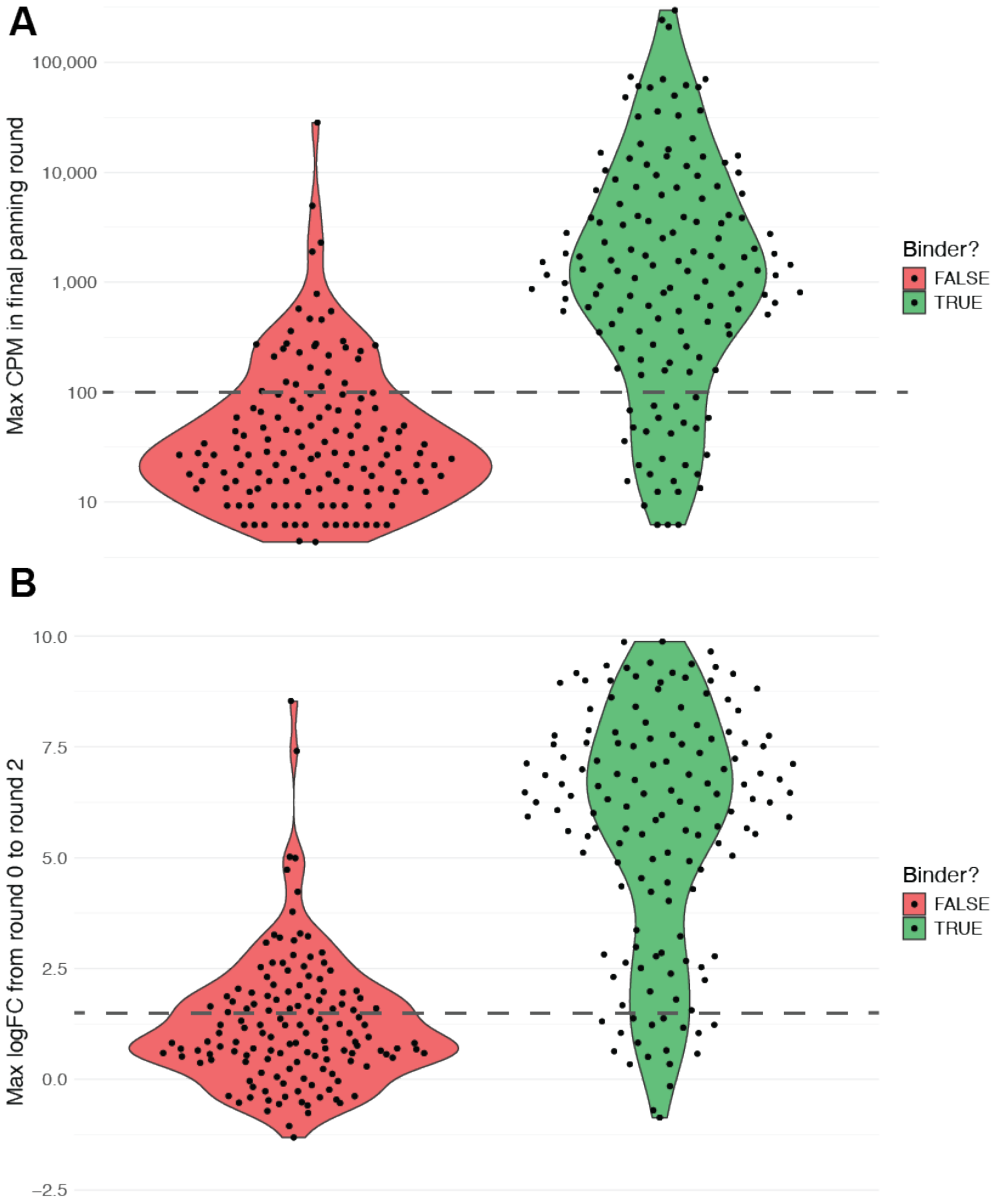


**Fig. 2. Relationship of nanobody binding to abundance and enrichment. (A)** Violin plots comparing the normalised counts per million (CPM) of nanobodies determined to be binders (green) or not (red) via either ELISA or BLI. A dotted line represents the empirically-derived cutoff of 100 CPM that was chosen as a default threshold to filter enriched sequences for candidate binders. **(B)** Violin plots comparing the enrichment, measured by $\log_2$ fold change (logFC) between round 0 and round 2 counts, of nanobodies determined to be binders (green) or not (red) via either ELISA or BLI. The *alpseq* pipeline implements a default threshold of logFC > 1.5 (dotted line) to prioritise enriched sequences among candidate binders.

#### Supplementary Note 3: Analysis of Hanke *et al.* (2020) dataset

The full interactive alpseq report is available here, and a static image of the interactive MDS plot/table is shown in Figure 3.

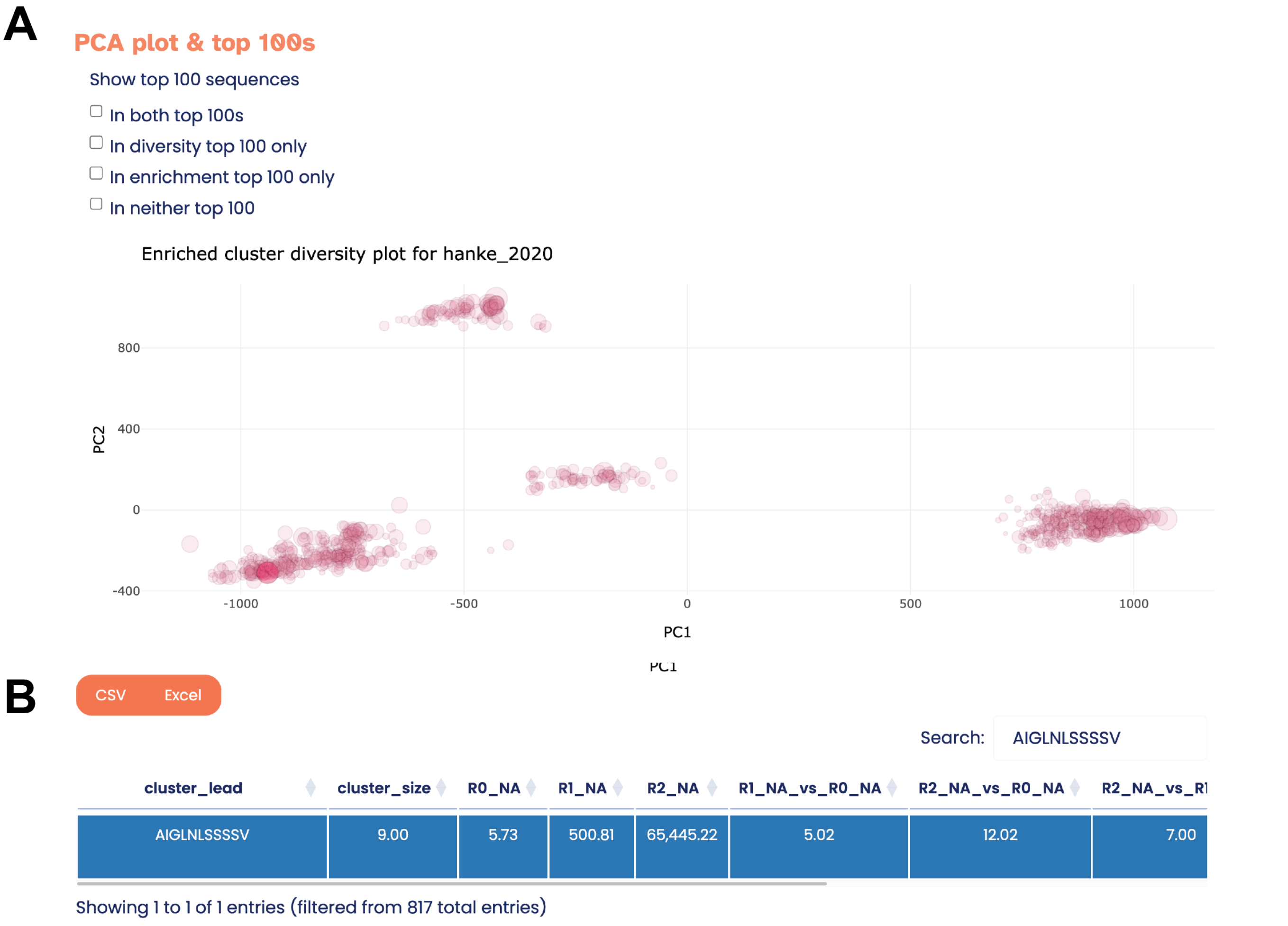


| cluster_lead | cluster_size | R0_NA | R1_NA | R2_NA | R1_NA_vs_R0_NA | R2_NA_vs_R0_NA | R2_NA_vs_R1 |
|---|---|---|---|---|---|---|---|
| AIGLNLSSSSV | 9.00 | 5.73 | 500.81 | 65,445.22 | 5.02 | 12.02 | 7.00 |

**Fig. 3. Enriched nanobody clusters from Hanke et al. dataset (A)** MDS plot of the sequence diversity landscape, showing how different nanobody clusters relate to one another based on their CDR3 sequences. The cluster of the Ty1 nanobody described in Hanke et al. (2020) is highlighted in red. **(B)** Table showing the CDR3, counts and logFC of the Ty1 nanobody cluster.

#### Supplementary Note 4: Analysis of Hanke *et al.* (2022) and Tsuruta *et al.* data

Selected results from our analyses of these public datasets are presented in Supplementary Figure 4 below. For the Tsuruta *et al.* dataset, we processed the raw data using *alpseq* and compared the cluster diversity plots for a few different pans. We observe that in the control and wild type samples, we don't see clear clusters (top row, Supplementary Figure 4A), but that these are present for the mutant samples (bottom row, Supplementary Figure 4A). For the Hanke *et al.* (2022) dataset, we processed the raw data using *alpseq* and compared the overlap between the three different pans in that study (Supplementary Figure 4B). We observed that, as expected based on the depletion of RBD binders in the competitive spike/RBD pan, there is no overlap between the enriched clones in those two pans.

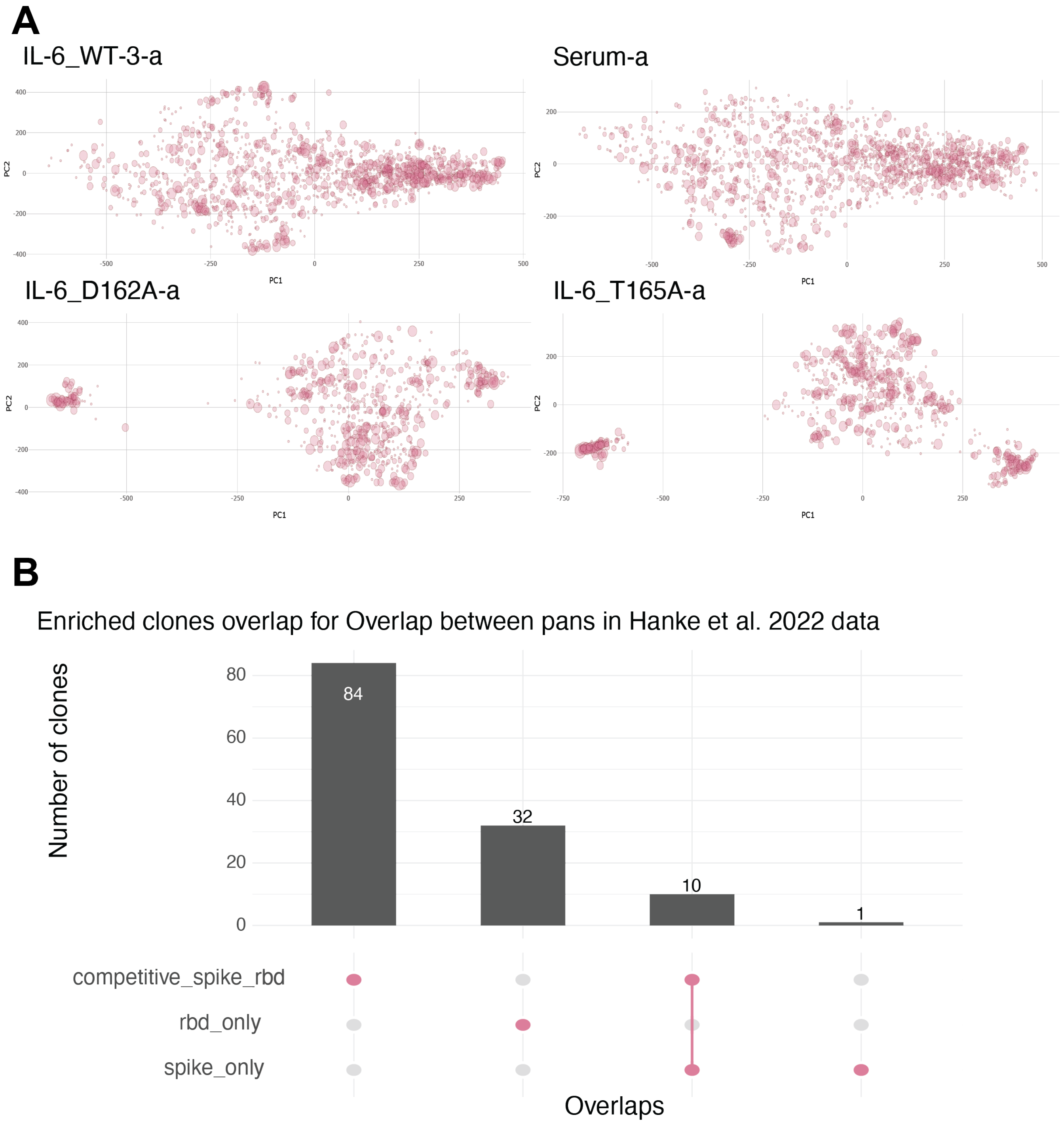


**Fig. 4. Further analyses of public datasets (A)** MDS plots of the sequence diversity landscape in the Tsuruta *et al.* data, showing how different nanobody clusters relate to one another based on their CDR3 sequences. Four different pans are chosen as representative examples. **(B)** UpSet plot of the overlap between the three pans in the Hanke *et al.* (2022) data.
